# GDP to GTP exchange on the microtubule end can contribute to the frequency of catastrophe

**DOI:** 10.1101/045765

**Authors:** Felipe-Andrés Piedra, Tae Kim, Emily S. Garza, Elisabeth A. Geyer, Alexander Burns, Xuecheng Ye, Luke M. Rice

## Abstract

Microtubules are dynamic polymers of αβ-tubulin that have essential roles in chromosome segregation and organizing the cytoplasm. Catastrophe – the switch from growing to shrinking – occurs when a microtubule loses its stabilizing GTP cap. Recent evidence indicates that the nucleotide on the microtubule end controls how tightly an incoming subunit will be bound (trans-acting GTP), but most current models do not incorporate this information. We implemented transacting GTP into a computational model for microtubule dynamics. In simulations, growing microtubules often exposed terminal GDP-bound subunits without undergoing catastrophe. Transient GDP exposure on the growing plus end slowed elongation by reducing the number of favorable binding sites on the microtubule end. Slower elongation led to erosion of the GTP cap and an increase in the frequency of catastrophe. Allowing GDP to GTP exchange on terminal subunits in simulations mitigated these effects. Using mutant αβ-tubulin or modified GTP, we showed experimentally that a more readily exchangeable nucleotide led to less frequent catastrophe. Current models for microtubule dynamics do not account for GDP to GTP exchange on the growing microtubule end, so our findings provide a new way of thinking about the molecular events that initiate catastrophe.

## INTRODUCTION

Microtubules (MTs) are hollow, cylindrical polymers of αβ-tubulin that have essential roles in organizing the cytoplasm and segregating chromosomes during cell division (reviewed in (Desai and Mitchison, 1997; Howard and Hyman, 2003)). Microtubules switch between phases of growing and shrinking, a hallmark property called dynamic instability that is directly targeted by some anti-mitotic drugs. Dynamic instability results from the biochemical and structural properties of the polymerizing αβ-tubulin subunits and how they interact with the microtubule lattice. Unpolymerized GTP-bound αβ-tubulin subunits assemble into microtubules, where their GTPase activity is greatly accelerated (Nogales *et al*., 1998). This assembly-dependent GTPase activity results in a ‘stabilizing cap’ of GTP-or GDP.Pi-bound αβ-tubulin at the end of growing microtubules. Loss of this stabilizing cap triggers catastrophe, the switch from growing to shrinking, by exposing the more labile GDP-lattice.

The microtubule-stabilizing role of GTP is widely appreciated, but there are conflicting views about the molecular mechanisms by which GTP stabilizes microtubules. Indeed, recent years have seen a change in thinking about the role of nucleotide in microtubule assembly and dynamics. Initially, a cis-acting view of nucleotide action dominated (Melki *et al*., 1989; Wang and Nogales, 2005; Nogales and Wang, 2006). This cis-acting view postulated that nucleotide state dictated the conformation of unpolymerized αβ-tubulin, with GTP favoring a straighter, microtubule-compatible conformation and GDP favoring a curved, microtubule incompatible conformation. This cis-acting nucleotide provided an appealing structural rationale to explain why microtubule polymerization requires GTP and why GTP hydrolysis leads to catastrophe. However, numerous biochemical and structural studies have failed to detect nucleotide-state-dependent conformational changes in unpolymerized αβ-tubulin that the cis-acting model predicts should occur (Honnappa *et al*., 2003; Rice *et al*., 2008; Barbier *et al*., 2010; Nawrotek *et al*., 2011; Ayaz *et al*., 2012; Pecqueur *et al*., 2012). Based on these and other studies, an alternative model for the role of GTP has gained favor(Buey *et al*., 2006; Rice *et al*., 2008). This ‘lattice’ or trans-acting model postulates that the effect of nucleotide state acts *across* the longitudinal interface, on a different αβ-tubulin than the one to which the nucleotide is bound. In a trans-acting model, the nucleotide does not affect the conformation/curvature of the unpolymerized αβ-tubulin to which it is bound. The difference between cis and trans models is most apparent when considering the terminal αβ-tubulin subunits at the growing end of the microtubule (Fig. 1): in a cis-acting model the nucleotide bound to the terminal subunit controls the strength of lattice contacts, but in a trans-acting model it is the nucleotide *underneath* (i.e. bound to a different αβ-tubulin) the terminal subunit that controls how tightly that terminal subunit associates with the lattice.

**Fig. 1.**
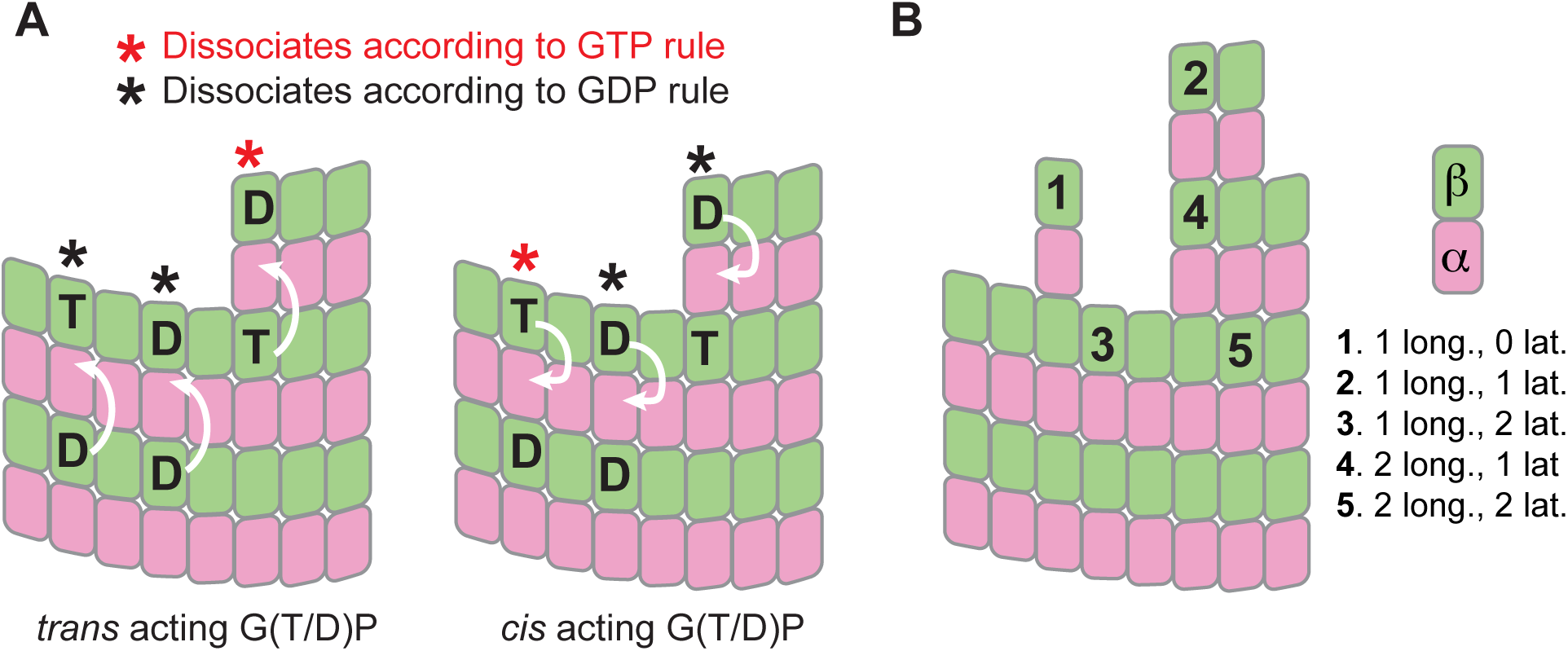
Cis and trans acting nucleotide in microtubule dynamics. (A) In the trans-acting model (left), the effect of nucleotide state is relayed across the longitudinal interface (cartooned by white arrows pointing from one αβ-tubulin to the one above). In the cis-acting model (right), the effect of nucleotide state is expressed on the subunit to which the nucleotide is bound (cartooned by white arrows pointing inside the same αβ-tubulin). ‘T’ and ‘D’ indicate nucleotide state, and red and black ‘*’ indicate which subunits make tighter lattice contacts. These different models made different predictions for how tightly terminal subunits will be bound. (B) In our model, the strength with which a given subunit is bound is determined by how many longitudinal and lateral interactions it makes, with more interactions giving tighter binding and slower dissociation. The effect of GDP is implemented as a multiplicative correction X such that subunits dissociating according to the ‘GDP rule’ will dissociate X-fold faster than their ‘GTP rule’ equivalents.

Most models for microtubule dynamics do not incorporate trans-acting nucleotide because these models were formulated when the exchangeable nucleotide was though to act in cis. Indeed, early models from Chen and Hill (Chen and Hill, 1983; 1985), later models from Martin and Bayley (Bayley *et al*., 1989; 1990; Martin *et al*., 1993), and more recent models from Odde and colleagues (VanBuren *et al*., 2002; 2005; Coombes *et al*., 2013)all share some version of cis-acting nucleotide. A more recent model(Margolin *et al*., 2012) began to explore how trans-acting nucleotide might affect polymerization dynamics but also invoked a protofilament cracking reaction that is difficult to validate experimentally. In the present study we sought to investigate potential functional consequences of trans-acting GTP by incorporating it into a computational model for microtubule dynamics, using only parameters that should be amenable to experimental perturbation. Following an approach that has been adopted by a number of other groups, we developed a Monte Carlo based algorithm that translates assumed biochemical properties of αβ-tubulin subunits into predictions of microtubule dynamics by simulating biochemical reactions – association, dissociation, and GTP hydrolysis – one at a time. For simplicity, we have not incorporated assumptions about αβ-tubulin mechanochemistry as has been done in other models (VanBuren *et al*., 2002; Coombes *et al*., 2013). Whether they incorporate mechanochemistry or not, these kinds of simulations provide a link between subunit biochemistry and the more ‘mesoscale’ parameters of microtubule dynamics: growing and shrinking rates, and the frequencies of catastrophe and rescue. Besides incorporating a trans-acting role for nucleotide, our model is conceptually most similar to an earlier model developed using a cis-acting mechanism for GTP/GDP (VanBuren *et al*., 2002) (Sup. Fig. 1).

Our simulations with trans-acting nucleotide revealed that transient exposure of GDP-bound αβ-tubulin on the microtubule plus end caused a slowing of elongation during which the stabilizing GTP cap eroded, increasing the propensity for catastrophe. This GTPase-dependent slowdown is not a unique consequence of trans-acting GTP, because we also observed it in a model using cis-acting nucleotide. We hypothesized that if GDP exposure led to catastrophe by reducing the rate of microtubule elongation, then GDP to GTP exchange on terminal subunits might mitigate the catastrophe-promoting effect of terminal GDP exposure. Our model calculations support this idea by showing that increased rates of GDP to GTP exchange on terminal subunits reduce the predicted frequency of catastrophe. We also performed experiments to perturb nucleotide-binding affinity (using mutant αβ-tubulin or modified nucleotides) with the goal of selectively increasing the rate of GDP to GTP exchange, observing that weaker nucleotide binding led to less frequent catastrophe. Thus, our simulations and experiments together support a model in which the rate of terminal nucleotide exchange can affect the frequency of catastrophe. These findings provide new ways of thinking about the molecular events that lead to catastrophe.

## RESULTS

We developed a kinetic Monte Carlo model to simulate MT polymerization dynamics one biochemical event (association, dissociation, GTP hydrolysis) at a time (part of the model has been described in (Ayaz *et al*., 2014)). Our model uses a two-dimensional representation of the microtubule lattice to capture different kinds of binding environments (Fig. 1A); it was inspired by one previously developed by Odde and Cassimeris (VanBuren *et al*., 2002). Our algorithm is constructed in a way that makes it easy to implement different biochemical assumptions. Here, we use our simulator with two different versions of the biochemical ‘rules’ governing polymerization and depolymerization: one set implements cis-acting GTP (Fig. 1B) as has been done previously (VanBuren *et al*., 2002)(Sup. Fig. 1), and the other set implements transacting GTP (Fig. 1B). For simplicity, we adopted a minimal parameterization that does not attempt to describe ‘mechanical’ properties of αβ-tubulin and microtubules such as spring-like conformational strain. This strain is thought to increase as αβ-tubulin changes from its curved, unpolymerized conformation to the straight conformation found in the polymer.

To obtain model parameters that could recapitulate MT elongation and shrinking rates and approximate the frequency of catastrophe, we followed the divide and conquer approach outlined in (VanBuren *et al*., 2002)(Methods). To more easily compare our model parameters to prior work, we trained our model on the same benchmark observations that have been used in other computational studies (Walker *et al*., 1988). First, we used ‘GTP only’ simulations to search for parameters that recapitulated MT growth rates over a range of αβ-tubulin concentrations (Walker *et al*., 1988) (Fig. 2A). With those parameters fixed, we optimized the weakening effect of GDP on the longitudinal interface by tuning it to make ‘all GDP’ microtubules depolymerize at the observed average rate of rapid, post-catastrophe shrinking. Because the growing and shrinking are taken to be ‘all GTP’ and ‘all GDP’ behaviors, the parameters that determine growing and shrinking rates are shared between simulations implementing cis-and trans-acting nucleotide. Finally, keeping the prior parameters fixed, we separately optimized the GTPase rate constant to obtain the observed catastrophe frequency at 10 |iM αβ-tubulin for cis and trans models (Fig. 2B,C). The parameters we obtained are comparable but not identical to those identified previously for the same data (VanBuren *et al*., 2002), validating our implementation. Compared to the cis-acting rules, the transacting ones require a slower GTPase rate to achieve the same frequency of catastrophe. Like the earlier study (VanBuren *et al*., 2002) that inspired ours, our model predicts a concentration-dependence of catastrophe that is steeper than the one measured in vitro, and does not recapitulate ‘age-dependent’ catastrophe. Nevertheless, our model provides a useful framework for investigating the possible effects of different biochemical assumptions.

**Fig. 2.**
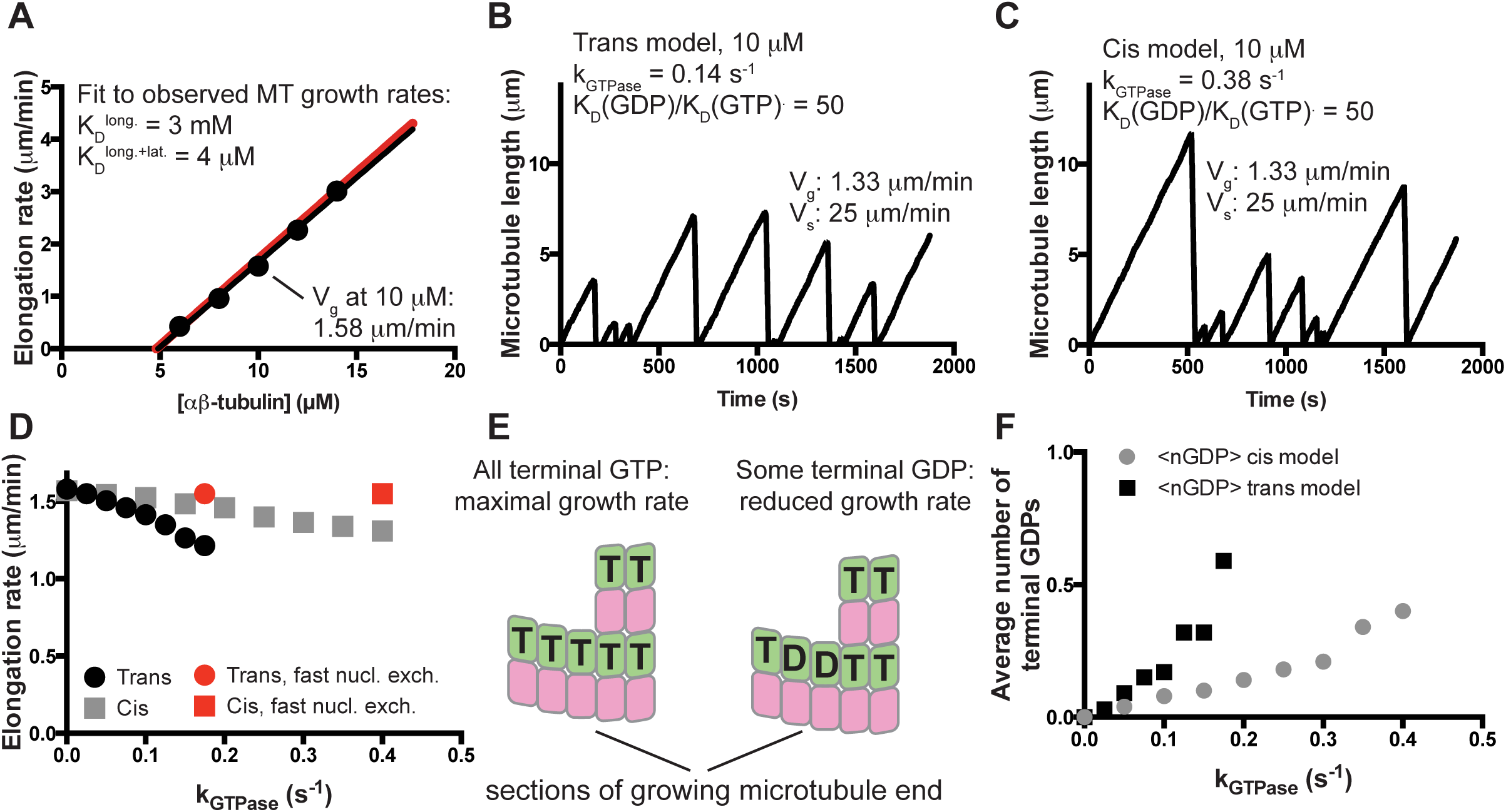
Kinetic Monte Carlo simulations of microtubule dynamics. (A) Simulations with ‘GTP only’ parameters recapitulate experimentally observed microtubule elongation rates over a range of mammalian αβ-tubulin concentrations (black circles: simulated elongation rates; black line: linear fit to the simulated elongation rates; red line: concentration-dependent elongation rates from (Walker *et al*., 1988)). (B) After optimizing the GTPase rate constant and the affinity weakening effect of GDP, the ‘trans’ model can recapitulate the experimentally observed frequency of catastrophe and rate of shrinking at 10 μM αβ-tubulin. Note that the elongation rate obtained here (1.33 μm/min) is slower than in the GTP only simulations (1.58 μm/min). (C) Same as B but using a version of cis-acting GTP (Sup. Fig. 1). Compared to the trans model, a faster rate of GTPase activity was required to obtain the measured frequency of catastrophe. A similar GTPase-induced ‘slowdown’ in elongation was observed. (D) Simulations show that for both trans and cis models the magnitude of the GTPase-induced slowdown increases with the GTPase rate. Red points indicate growth rates in simulations where terminal GDP to GTP exchange was instantaneous. (E) Exposure of terminal GDP-bound αβ-tubulins might provide an explanation for the GTPase-induced decrease in elongation rates. (F) In both trans and cis simulations, microtubules expose about 1 terminal GDP subunit every three end configurations sampled during a growth phase.

When a non-zero GTPase rate constant was used in simulations, the elongation rates decreased from their ‘GTP only’ values for both cis and trans models (e.g. by 15% in Fig. 2B,C). To our knowledge, such a GTPase-induced ‘slowdown’ has not previously been reported. Simulations using a range of GTPase rate constants revealed that for both cis and trans models the rate of microtubule elongation decreased monotonically from its ‘GTP only’ value as the GTPase rate increased (Fig. 2D). We speculated that the GTPase-induced slowdown occurred because GDP-tubulin was being exposed on the microtubule end, creating low affinity binding sites that antagonized elongation enough to reduce average growth rates without triggering catastrophe (Fig. 2E). If true, then the microtubules in our simulations should have increasing amounts of GDP-bound terminal subunits as the GTPase rate constant is increased. Counting the number of terminal GDP-bound subunits during growth revealed that the simulated microtubules do indeed expose GDP-bound terminal subunits, and that the frequency of exposure increases with increasing GTPase rate (Fig. 2F). Using the parameters from Fig. 2B,C, ‘cis’ and ‘trans’ microtubules on average each expose 1 terminal GDP subunit ~33% of the time during a growth phase. In other words, on average every third configuration sampled during growth contained one terminal GDP. To confirm that the GTPase-induced slowdown did in fact result from exposing GDP-bound terminal subunits, we modified our simulation code to instantaneously convert terminal GDP-tubulin subunits back to the GTP state. This modification resulted in elongation rates indistinguishable from those obtained in the absence of GTPase (red points in Fig. 2D).

The experiments described above indicate (i) that GDP-bound terminal subunits can be exposed on the end of growing microtubules, (ii) that this exposure slows elongation rates, and (iii) that the GTPase-induced slowdown is abrogated when GDP to GTP exchange on terminal subunits occurs instantaneously. We wondered if a finite rate of GDP to GTP exchange on terminal subunits might affect the frequency of catastrophe by modulating the GTPase-induced slowdown in our simulations (Fig. 3A). To test this, we added a terminal GDP to GTP exchange reaction to our algorithm (Fig. 3B), assuming that GDP release was rate-limiting for exchange. The rate of terminal GDP to GTP exchange markedly affected the predicted frequency of catastrophe in simulations with faster rates reducing the frequency of catastrophe (Fig. 3B). At very fast rates of terminal nucleotide exchange the catastrophe frequency approached zero in the trans model, and at very slow rates of exchange the catastrophe frequency approached a maximum value. At intermediate rates of nucleotide exchange, the predicted catastrophe frequency depends sensitively on the rate of exchange (Fig. 3B).

**Fig. 3.**
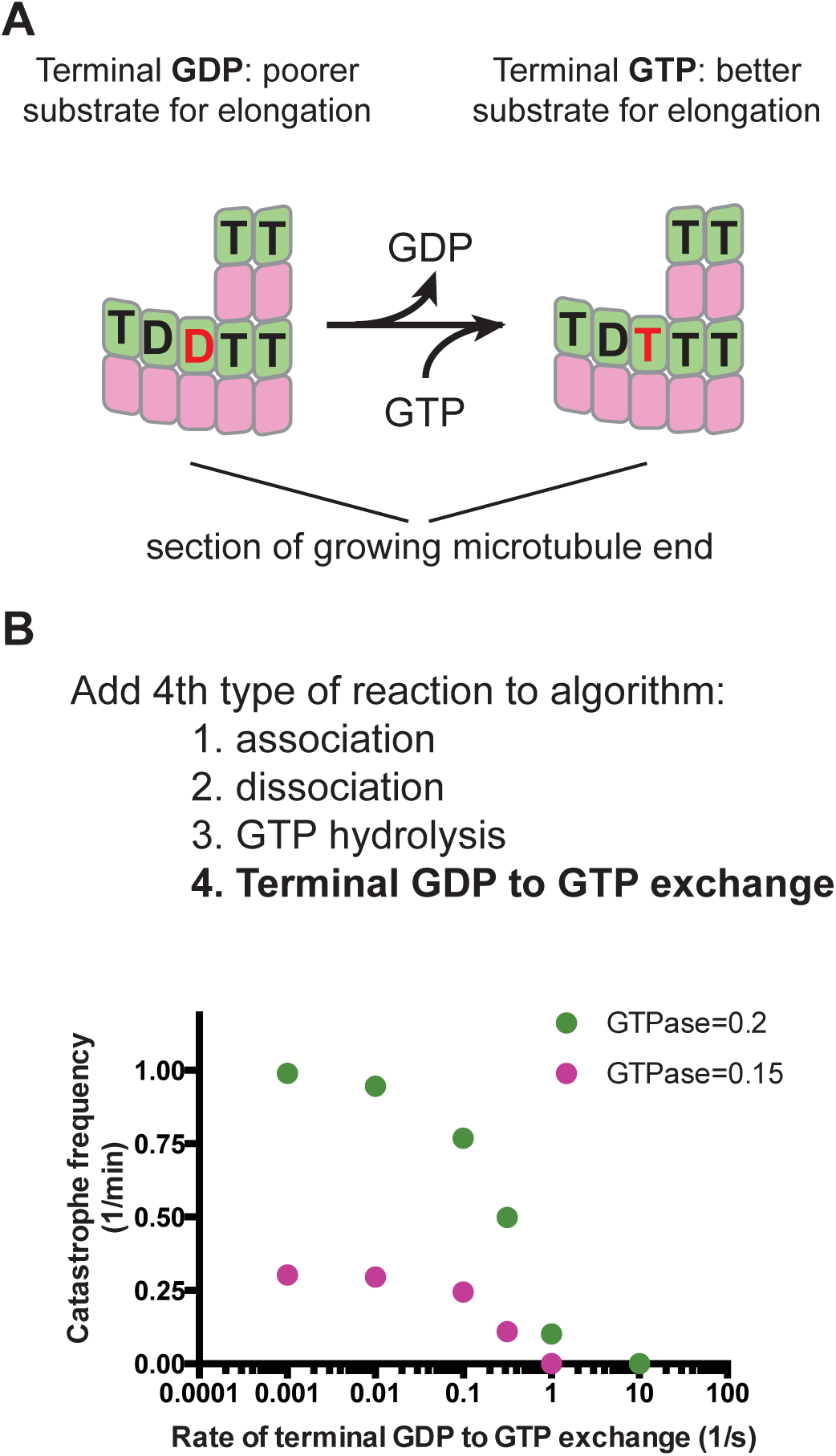
A possible role for terminal GDP to GTP exchange in microtubule catastrophe. (A) Cartoon illustrating GDP to GTP exchange on a terminal subunit. (B) To investigate in simulations the possible relevance of terminal GDP to GTP exchange, we added it as another possible event in our simulation algorithm. Varying the rate of terminal GDP to GTP exchange in simulations with two different GTPase rate constants (green: 0.2 s^−1^, pink: 0.15 s^−1^) yielded a marked effect on the predicted frequency of catastrophe, with faster rates of exchange giving less frequent catastrophe.

Observing a GDP-induced ‘slowdown’ in simulations led us to wonder if actual microtubules regularly expose GDP-bound subunits on their growing plus ends, and if terminal nucleotide exchange might affect their frequency of catastrophe. In what follows, we use two complementary perturbations of nucleotide binding to explore if GDP to GTP exchange on the growing microtubule end affects the frequency of catastrophe in vitro, using yeast αβ-tubulin as a model system because it has allowed in vitro studies using site-directed mutants (Davis *et al*., 1994; Sage *et al*., 1995; Dougherty *et al*., 1998; 2001; Gupta *et al*., 2002; Uchimura *et al*., 2010; Johnson *et al*., 2011; Geyer *et al*., 2015).

An αβ-tubulin mutant that exchanges nucleotide faster but that is otherwise normal could provide a way to test the prediction that faster terminal exchange reduces the frequency of catastrophe. Although directly measuring the rate of nucleotide exchange on the MT end is not yet possible, we reasoned that faster exchange would likely result from a decrease in αβ-tubulin GTP/GDP binding affinity, which is easily measured. To affect nucleotide binding as selectively as possible, we sought to mutate GTP-interacting residues that do not also participate in polymerization contacts or GTPase activity. The conserved β-tubulin residue C12 fits these criteria: the sidechain packs underneath the guanosine base of the exchangeable nucleotide, does not contact neighboring αβ-tubulins, and has not been implicated in catalysis (Fig. 4A). We therefore overexpressed and purified (Johnson *et al*., 2011) C12A yeast αβ-tubulin in order to study its GTP-binding and polymerization dynamics.

**Fig. 4.**
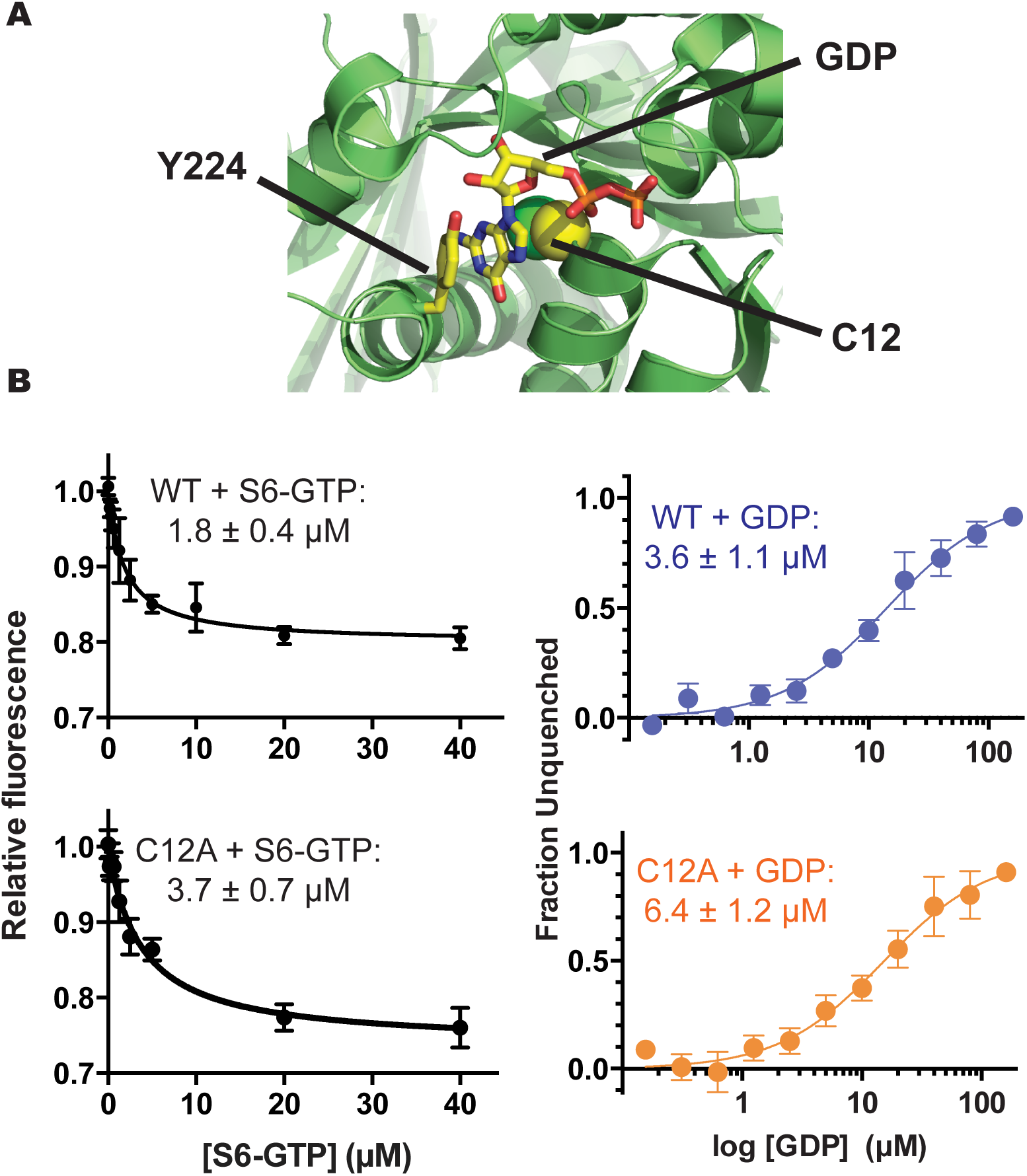
Nucleotide-binding properties of β:C12A αβ-tubulin. (A) β:C12 (sidechain represented as spheres) is a conserved residue that contacts the exchangeable GTP but that does not participate in polymerization contacts. The view, which is approximately of the plus-end of β-tubulin, shows part of the surface that would be contacted by another αβ-tubulin; Y224 shields the guanosine from polymerization contacts by stacking on top of it. PDB 4I4T (Prota *et al*., 2013) was used for this illustration. (B) The β:C12A mutation reduces the nucleotide-binding affinity of αβ-tubulin. (left panels) The affinities of wild-type and β:C12A αβ-tubulin for S^6^-GTP were measured by monitoring quenching of αβ-tubulin intrinsic fluorescence by S^6^-GTP. Each data point represents the mean fraction of signal quenched at a particular concentration of S^6^-GTP (n = 4; Error bars represent the standard error of the mean). (right panels) The affinities of wild-type and β:C12A αβ-tubulin for GDP were measured using a fluorescence unquenching assay. Each data point represents the mean fraction of signal restored at a particular concentration of GDP (n = 6; Error bars represent the standard error of the mean).

We first measured the nucleotide binding affinity of C12A yeast αβ-tubulin using a fluorescence-quenching assay (Fishback and Yarbrough, 1984; Amayed *et al*., 2000). This assay takes advantage of the fact that 6-thio-GTP (S^6^-GTP) quenches the intrinsic fluorescence of αβ-tubulin when bound. We first determined the affinity of wildtype and C12A αβ-tubulin for S^6^-GTP, obtaining values of 1.8 and 3.7 μM, respectively (Fig. 4B). We next used a competition assay to obtain their affinity for GDP. Wildtype αβ-tubulin binds GDP with K_D_ = 3.6 μM; C12A binds GDP less tightly, with K_D_ = 6.4 μM (Fig. 4B). A prior measurement of GTP binding to yeast αβ-tubulin showed higher affinity (~50 nM; (Davis *et al*., 1993)); this discrepancy may be explained by the glycerol-free conditions we used (glycerol has been observed to increase the nucleotide binding affinity of αβ-tubulin (Yarbrough and Fishback, 1985)). Whatever the reason, our measurements showed that the β:C12A mutation caused a decrease in the affinity of GTP/GDP binding.

To determine if the β:C12A mutation also affected the apparent biochemistry of microtubule elongation and/or the frequency of catastrophe, we used time-lapse differential interference contrast microscopy to measure the polymerization dynamics of wild type and β:C12A αβ-tubulin. β:C12A and wild-type microtubules showed similar concentration-dependent growth rates. This similarity suggests that the mutation had little effect on the affinity of the interactions that drive elongation (estimated from extrapolation to the x-axis, 51 nM and 42 nM for wild-type and β:C12A, respectively) or on the apparent rate constant for subunit addition (estimated from the slope/concentration-dependence of elongation rates, 14.9 μM^−1^s^−1^ and 12.2 μM^−1^s^−1^ for wild-type and β:C12A, respectively) (Fig. 5A). Despite the somewhat slower growth rates at a given concentration, β:C12A MTs underwent catastrophe substantially less frequently than wildtype, showing a 6-fold difference at concentrations around 0.3 μM (Fig. 5B).

**Fig. 5.**
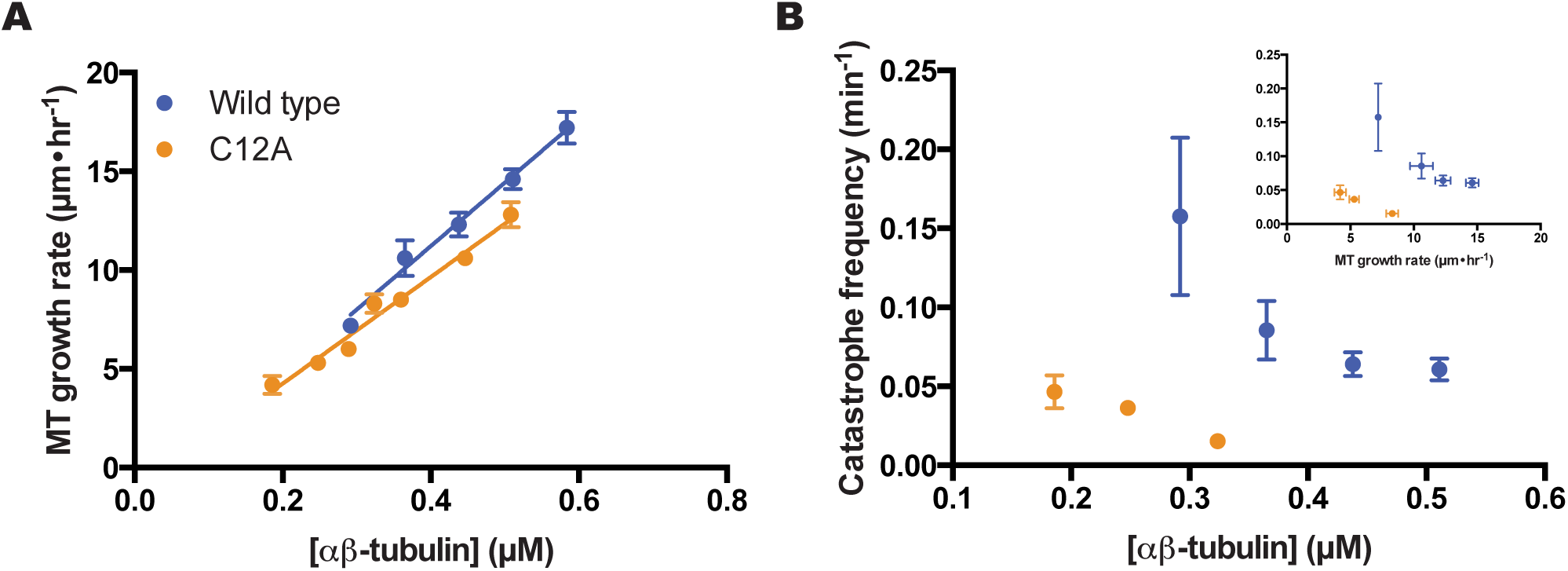
β:C12A microtubules undergo catastrophe less frequently than wild-type. (A) β:C12A microtubules show comparable concentration-dependent elongation rates as wild-type (n=4 microtubules per data point, error bars show the standard deviation). Parameters (slope; y-intercept) from the linear regressions are: wild-type (14.9 +/- 0.9 μM^−1^s^−1^; −0.9 +/- 0.4 s^−1^) and β:C12A (12.2 +/- 0.3 μM^−1^s^1^; −0.5 +/- 0.3 s^−1^). (B) β:C12Amicrotubules undergo catastrophe ~7-fold less frequently than wild-type. Inset: A plot of catastrophe frequency vs. MT growth velocity. The small changes in growth rate do not explain the change in catastrophe frequency. (Each data point represents the catastrophe frequency measured at a particular concentration of αβ-tubulin (from L to R for wild type (blue): n = 10, 21, 75, 75; from L to R for β:C12A (orange): n = 20, 90, 63); Error bars represent the standard deviation).

To further probe a potential link between the rate of terminal GDP to GTP exchange and the frequency of microtubule catastrophe, we sought an alternative way to perturb nucleotide binding affinity that would be free of the complications associated with mutations. Earlier work showed that deoxy analogs of GTP promote MT assembly and bind less tightly to the exchangeable site (Hamel *et al*., 1984). Structures of αβ-tubulin (e.g. (Prota *et al*., 2013) for a recent high resolution example) further show that the 2’ hydroxyl group of the guanine nucleotide does not make polymerization contacts (Fig. 6A). Thus, measuring microtubule dynamics in the presence of dGTP should provide an alternative to site-directed mutants. Using the competition assay described above, we determined that yeast αβ-tubulin binds dGDP about 2-fold less tightly than GDP (7.3 μM vs 3.6 μM; Fig. 6B), a change in affinity consistent with prior measurements using mammalian αβ-tubulin (Hamel *et al*., 1984). Measured MT elongation rates changed little with dGTP compared to GTP (concentration-dependence for elongation rates: 15.8 μM^−1^s^−1^ vs 14.9 μM^−1^s^−1^ for wild-type; extrapolation to the x-axis: 30 nM vs 61 for wild-type) (Fig. 6C), indicating that the removal of the 2’-OH did not substantially perturb the strength of lattice contacts. Even accounting for the slightly faster elongation, however, the catastrophe frequency decreased markedly in the presence of the weaker binding dGTP (over 4-fold decrease at 0.3 μM) (Fig. 6D).

**Fig. 6.**
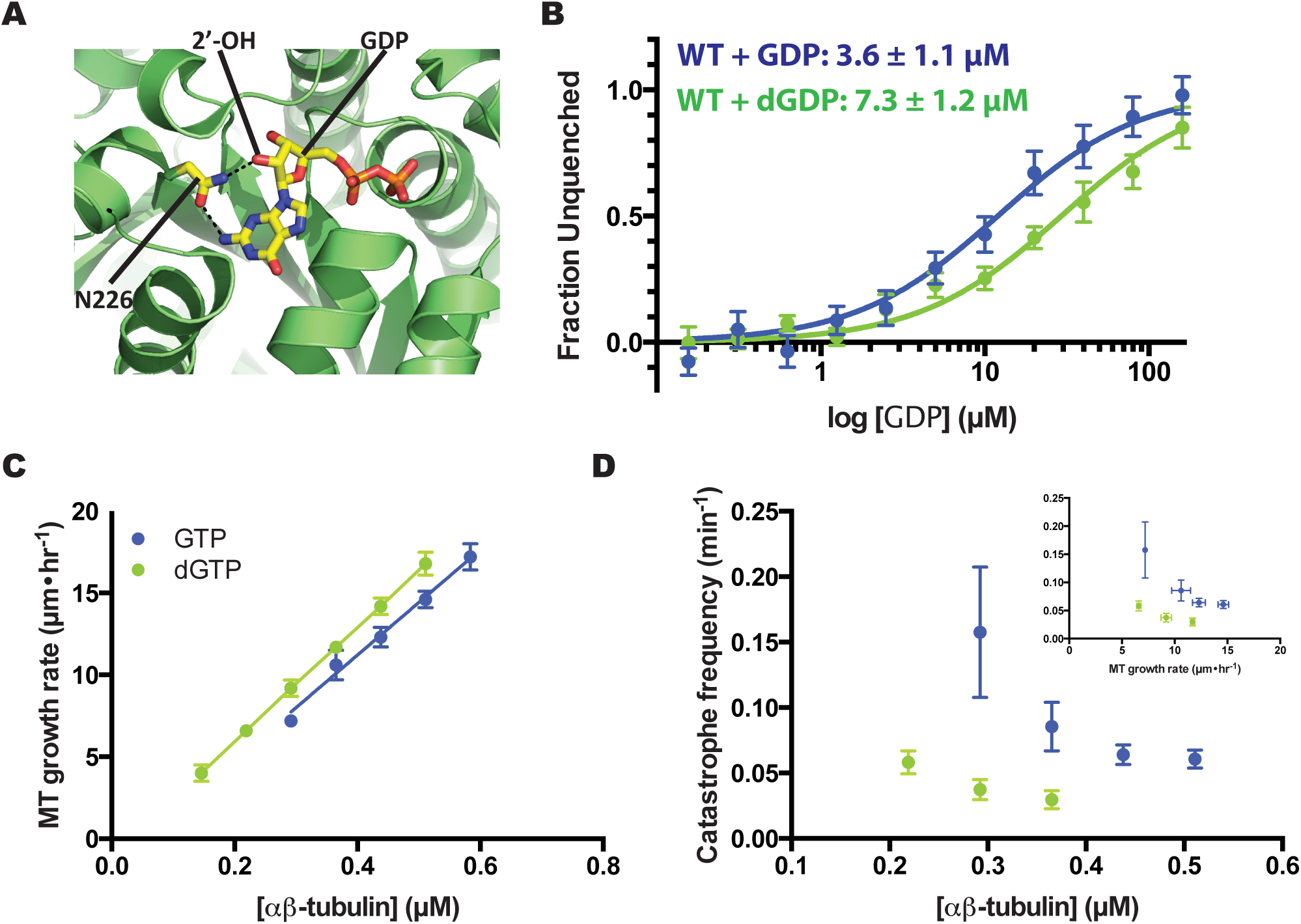
αβ-tubulin binds less tightly to, and undergoes catastrophe less frequently with, 2’-deoxy guanine nucleotides. (A) (left) The 2’ hydroxyl of GTP, which is absent in dGTP, makes a hydrogen-bond with a conserved asparagine (N202). PDB 4I4T (Prota *et al*., 2013) was used for this illustration. (right) The affinities of yeast αβ-tubulin for GDP and dGDP were measured using a fluorescence-based competition assay (see Methods). αβ-tubulin binds to dGDP about 2-fold less tightly than to GDP. (Each data point represents the mean fraction of signal restored measured at a particular concentration of GDP or dGDP (n = 6 for GDP (blue); n = 4 for dGDP (green)); Error bars represent the standard error of the mean). (B) MTs polymerizing with dGTP show comparable concentration-dependent elongation rates to those growing with GTP MTs (n=4 MTs per data point, error bars show the standard deviation). Parameters (slope; y-intercept) from the linear regression for elongation in the presence of dGTP are: 15.8 +/- 0.06 μM^−1^s^−1^; - 0.5 +/- 0.02 s^−1^). (C) MTs assembled with dGTP undergo catastrophe ~4-fold less frequently than with GTP. Inset: A plot of catastrophe frequency vs. MT growth velocity shows that the small increase in growth rate cannot explain the change in catastrophe frequency. (Each data point represents the catastrophe frequency measured at a particular concentration of αβ-tubulin (from L to R for wild-type + GTP (blue): n = 10, 21, 75, 75; from L to R for wild-type + dGTP (green): n = 43, 23, 18); Error bars represent the standard deviation).

A decreased frequency of catastrophe for two different perturbations that reduce nucleotide binding affinity is consistent with our proposal that the rate of terminal nucleotide exchange can contribute to the frequency of catastrophe. However, the experiments described above do not rule out an alternative mechanism in which reduced GTPase activity (as a result of the mutation and/or the use of dGTP) is responsible for the reduced frequency of catastrophe. We used two approaches to investigate the possibility that the β:C12A mutation or the use of dGTP affected GTPase activity. We first performed experiments with ^32^P-GTP to compare the nucleotide content of wild-type microtubules (polymerizing with GTP) to β:C12A microtubules (with GTP) and to wildtype microtubules polymerizing with dGTP. These data show that wild-type and β:C12A microtubules contain comparable amounts of GTP (50% and 48%, respectively), and that when polymerizing with dGTP wild-type microtubules contain less GTP (25%) (Fig. 7A). These data indicate that the β:C12A mutation has little if any effect on GTPase in the microtubule, and that dGTP appears to somewhat increase GTPase rates. These GTP contents are higher than expected, so we used Bim1 (Schwartz *et al*., 1997), an EB1 family protein that reports on features of the microtubule lattice thought to be related to nucleotide state (Maurer *et al*., 2012; Zhang *et al*., 2015), as an alternative readout. Wildtype and β:C12A microtubules have Bim1 caps of very similar size and maximal intensity, and Bim1 binds comparably to the non-cap lattice of wild-type and mutant microtubules (Fig. 7B,C). The lack of substantial differences in the binding of Bim1 to wild-type and β:C12A microtubules suggests that any differences in the distribution of GTP must be small. These data also argue against a potential allosteric effect of the mutation like the one we recently described for β:T238A αβ-tubulin (Geyer *et al*., 2015). In contrast, wild-type microtubules growing with dGTP show narrower and less intense Bim1 caps, which suggests that the dGTP is either more rapidly hydrolyzed or that it perturbs microtubule structure in a way that reduces Bim1 binding. Taken together, the ^32^P-GTP and Bim1 experiments argue against an alternative model in which a GTPase defect is responsible for the reduced catastrophe frequency of β:C12A or wild-type(dGTP) microtubules.

**Fig. 7.**
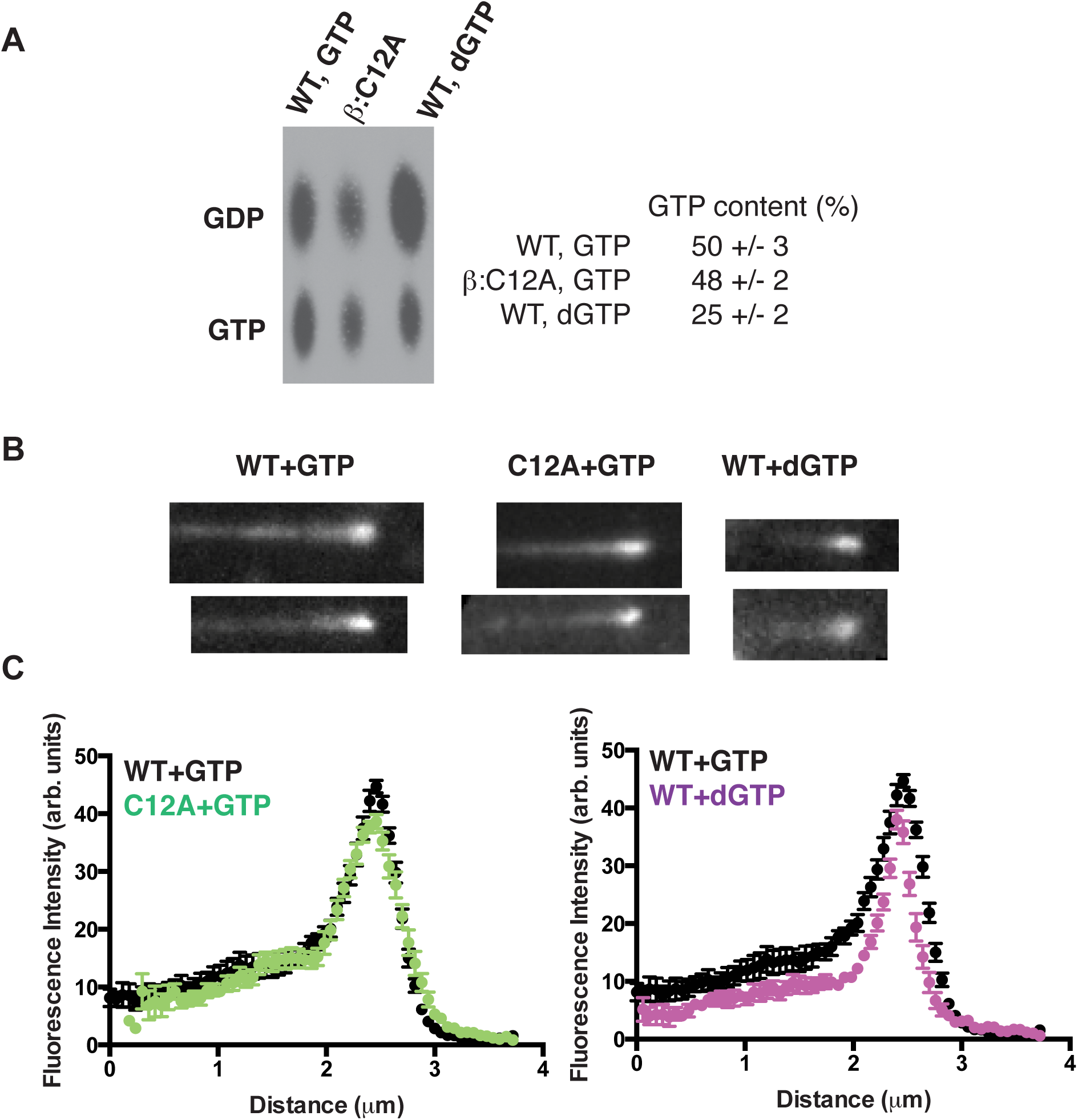
Exchangeable GTP content of microtubules probed with ^32^P-GTP and with an EB1 protein. (A) Image shows ^32^P nucleotide content of wild-type (grown with GTP or dGTP) or β:C12A mutant microtubules (grown with GTP) analyzed by TLC. Data from 4 independent trials are tabulated. Wild-type and β:C12A microtubules growing with GTP show comparable ^32^P-GTP content; wild-type microtubules growing with dGTP show reduced ^32^P-GTP content. (B) Representative images of microtubules growing in the presence of Bim1-GFP. Scale bar = 1 μm. (C) Plots of average Bim1-GFP intensity (n=10) on microtubules. Wild-type and β:C12A microtubules growing with GTP show very similar Bim1 comets, whereas wild-type microtubules growing with dGTP show a narrower comet.

To better understand how terminal GDP exposure could contribute to microtubule catastrophe and elongation rate, we examined simulated growth phases more closely (Fig. 8). We extracted the length of the microtubule (as determined by the total number of αβ-tubulin subunits that have been incorporated), the size of the GTP cap (number of incorporated subunits that are bound to GTP; the remainder are bound to GDP), and the number of GDP-bound subunits exposed on the growing end. A representative plot of simulated microtubule length vs. time is shown in Fig. 8A; the two boxed regions are shown with higher temporal and spatial resolution in Fig. 8B,C. The simulations show transient pauses (sometimes involving net loss of subunits) in the rate of microtubule elongation (top panels in Fig. 8B,C; the time at which pauses begin are indicated by grey arrows) that are associated with the exposure of a small number of terminal GDP-bound subunits (middle panels in Fig. 8B,C). The stabilizing GTP cap – which fluctuates around a steady-state value of ~200 GTP-bound αβ-tubulins under the conditions of our simulations – erodes as a consequence of the pausing. In our simulations the pausing and erosion of the GTP cap that are induced by exposure of terminal GDP-bound subunits can sometimes lead to catastrophe (black arrow in Fig. 8C), but not always. Indeed, elongating microtubules can ‘recover’ normal elongation rates and GTP caps by eliminating the terminal GDP-bound subunits (Fig. 8B,C). GDP-to-GTP exchange on terminal subunits is one of the biochemical pathways that allow recovery from pausing and its attendant erosion of the GTP cap.

**Fig. 8.**
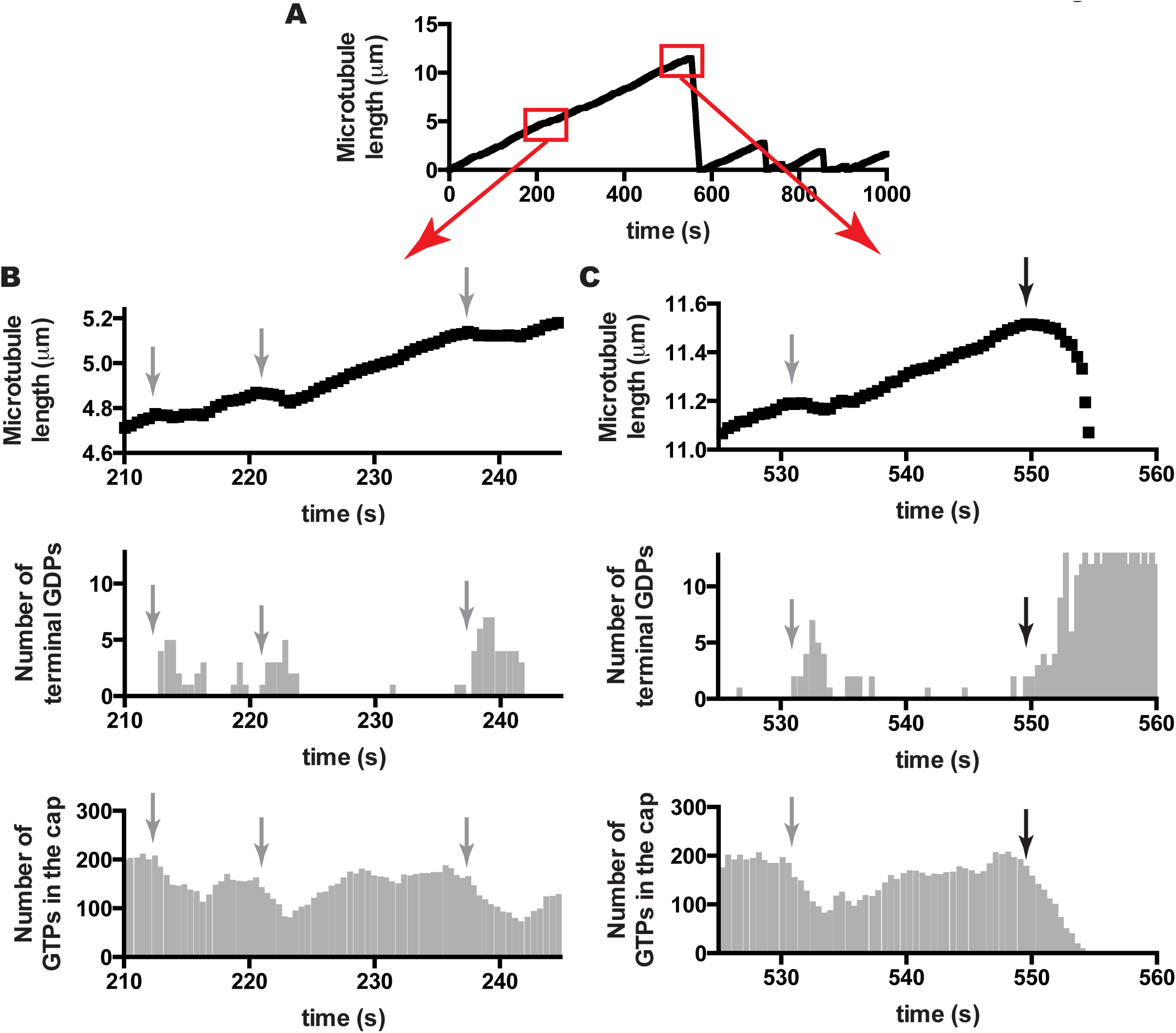
Insights into catastrophe from the model. (A) A length history plot from a simulation using model parameters (from Fig. 2B) and 10 uM αβ-tubulin. Boxed regions are shown in greater detail in the subsequent panels. (B) (top) Transient reductions in the elongation rate (indicated by grey arrows) are apparent when the growth phase is plotted with greater temporal and spatial resolution. (middle) The pausing occurs when GDP-bound subunits are exposed on the microtubule end, consistent with the model advanced in Fig. 2. (bottom) When microtubule elongation slows, the GTP cap erodes. Elimination/reduction of GDP-bound subunits on the microtubule end, which can occur through multiple mechanisms including terminal GDP to GTP exchange, restores more normal growth rates that allow the GTP cap to recover. The GTP cap fluctuates around its steady state value of about 190 under these conditions. Faster nucleotide exchange on the microtubule end contributes to increase the likelihood of these ‘mini-rescues’ that occur as a consequence of terminal GDP exposure. (C) (top) Catastrophes in our simulations are also preceded by a reduction in elongation rate (black arrow marks the onset of the slowdown). This slowing before catastrophe is reminiscent of that seen in a recent study of EB1. (middle) As with the transient pausing, the slowing before catastrophe is associated with the exposure of GDP-bound terminal αβ-tubulin subunits. (bottom) The disappearance of the GTP cap occurs over the course of several seconds. The microtubule can have all terminal subunits bound to GDP before the cap has been completely lost.

## DISCUSSION

We began this study using a computational model to investigate the consequences of trans-acting nucleotide for microtubule dynamics. Our simulations with trans-acting nucleotide revealed that GDP-terminated protofilaments were frequently exposed during sustained growth without necessarily causing catastrophe. This apparently contrasts with a common assumption associated with cis-acting GTP - that the exposure of GDP-bound αβ-tubulin ‘poisons’ the protofilament (VanBuren *et al*., 2005). Our computational simulations further revealed that transient exposure of GDP-bound αβ-tubulins on the microtubule end reduces the rate of elongation, and that this in turn erodes the GTP cap and increases the likelihood of MT catastrophe. GDP to GTP exchange on terminal GDP-bound subunits can modulate the frequency of catastrophe by mitigating the effects of GDP exposure on growing microtubule ends.

Current thinking about catastrophe is dominated by considerations about the size of the microtubule’s stabilizing GTP cap and the release of strain from the resulting GDP lattice (Bowne-Anderson *et al*., 2013; Maurer *et al*., 2014). However, the molecular events that cause loss of the cap during otherwise steady state growth continue to remain poorly understood (Gardner *et al*., 2013; Bowne-Anderson *et al*., 2015; Brouhard, 2015). GDP to GTP exchange on the growing microtubule end represents a simple biochemical mechanism that is spatially segregated from GTP hydrolysis and that provides a new way of thinking about how catastrophe might be initiated. That terminal nucleotide exchange could be relevant to catastrophe is not a new idea. Indeed, GDP to GTP exchange on the growing microtubule end figured in some of the earliest computational models for microtubule dynamics (Chen and Hill, 1983; 1985). Early in vitro studies also showed that the terminal nucleotide was exchangeable (Mitchison, 1993; Caplow and Shanks, 1995), and that modulating the amount of terminal GDP exposure could alter the frequency of catastrophe (Caplow and Shanks, 1995; Vandecandelaere *et al*., 1995). Our experiments with mutant αβ-tubulin and deoxy-GTP sought to perturb nucleotide binding without affecting other aspects of αβ-tubulin biochemistry. Those experiments showed that reduced GTP/GDP-binding affinity results in a lower frequency of microtubule catastrophe, providing support for the idea that the rate of GDP to GTP exchange on the growing microtubule end contributes to catastrophe. The GDP-exposure-induced slowing of elongation and ensuing erosion of the GTP cap that we observed in simulations is reminiscent of a recent study of the end-binding protein EB1, wherein microtubule growth was observed to slow down before the EB1-marked stabilizing cap began to shrink (Maurer *et al*., 2014). We speculate that exposure of terminal GDP-bound subunits contributes to that slowdown.

Our goal was to explore a simple model incorporating trans-acting nucleotide. A surprising observation from the simulations led to the ideas about terminal GDP exposure and the role of terminal GDP to GTP exchange. The minimal model that we developed does not capture the concentration-or age-dependence of microtubule catastrophe (Gardner *et al*., 2011b; 2013), and also does not attempt to model allosteric/mechanochemical properties that are likely to have important roles in catastrophe (VanBuren *et al*., 2005; Zanic *et al*., 2013; Geyer *et al*., 2015). It seems that to successfully recapitulate the full spectrum of catastrophe, and to place terminal nucleotide exchange into the broader context of allostery and mechanochemistry, future models will need to incorporate aspects from several current models: trans-acting nucleotide, GDP to GTP exchange on the microtubule end, multiple conformations and ‘spring-like’ mechanical properties of αβ-tubulin, and perhaps also neighbor-state specific rates of GTP hydrolysis and/or exchange. We expect that progress on this challenging goal will accelerate in light of recently developed methods that enable in vitro work with site-directed αβ-tubulin mutants(Georges *et al*., 2008; Johnson *et al*., 2011; Widlund *et al*., 2012; Minoura *et al*., 2013), and because experimental observations of catastrophe are being extended to higher spatial and temporal resolution (Gardner *et al*., 2011b; Maurer *et al*., 2014).

## MATERIALS AND METHODS

### Kinetic Modeling

We wrote a computer program to perform kinetic Monte Carlo simulations of MT dynamics, implementing an algorithm similar to one described previously (VanBuren *et al*., 2002). Part of our implementation has been described elsewhere (Ayaz *et al*., 2014). Briefly, the microtubule lattice is represented by a two dimensional grid to account for the different kinds of binding sites on the microtubule end, and with a staggered periodic boundary condition to mimic the cylindrical microtubule structure (Ayaz *et al*., 2014). A portion of the lattice was treated as permanently occupied to provide a seed for elongation. The program simulates MT dynamics one biochemical reaction (subunit addition or dissociation, GTP hydrolysis, or GDP to GTP exchange) at a time. Our parameterization assumes that the nucleotide (GTP or GDP) acts in trans to affect the strength of longitudinal contacts with GTP contacts stronger than GDP ones (Figure 1A, and see below); prior models have assumed cis-acting GTP (Bayley *et al*., 1989; VanBuren *et al*., 2002; 2005; Brun *et al*., 2009; Gardner *et al*., 2011a; Coombes *et al*., 2013) (but see (Margolin *et al*., 2012) for a notable exception). The rate of subunit addition into any available site is given by k_on_*[αβ-tubulin]. Occupied sites (excepting the seed) can dissociate at a rate given by k_on_*K_D_^site^ where K_D_^site^ is the affinity of interaction at a particular site, which is determined by the neighbor state (number and type of lattice contacts) and obtained from longitudinal and corner affinities through thermodynamic coupling (Erickson and Pantaloni, 1981; VanBuren *et al*., 2002). GTP hydrolysis is modeled for all nonterminal subunits with rate constant kGTPase. The weakening effect of GDP on the longitudinal interface is modeled as a multiplicative scaling of the longitudinal contribution – this is equivalent to a loss of favorable free energy of association. Because of the high concentration of GTP in our assays (1 mM), we assume that GDP release is rate limiting for GDP to GTP exchange. Accordingly we model exchange on terminal subunits with rate constant k_exchange_. Simulations begin with only 13 possible events (associations onto the end of each protofilament). Execution times for each event are determined by sampling a random number x between 0 and 1, and then calculating the time as -(1/*rate*)*ln(x), where *rate* gives the appropriate rate constant as described above. At each step the event with the shortest execution time is implemented, the simulation time is advanced accordingly, and the list of possible events and their associated rates is updated to account for changes in subunit neighbor state. To obtain the length (in μm) of a simulated microtubule at a given time, we divide the number of subunits by 1625 (one αβ-tubulin is 8 nm in length).

### Parameter optimization

To recapitulate experimentally observed, concentration-dependent rates of MT elongation, and as described previously (Ayaz *et al*., 2014), we performed a manual grid search using ‘GTP only’ (negligible GTPase rate constant) simulations to obtain longitudinal and corner (longitudinal+lateral) affinities, initially ignoring GDP to GTP exchange on the MT end. With these parameters fixed we next searched for the ‘GDP-weakening’ factor that would yield the experimentally observed concentration-independent rate of rapid microtubule shrinking. Finally, we searched for a GTPase rate constant that would give the appropriate frequency of catastrophe (Fig. 2B). This procedure essentially followed the approach outlined in (VanBuren *et al*., 2002), and yielded similar values for the parameters in common between the two models. For an assumed on-rate constant of 4x10^6^ M^-1^sec^-1^, we obtained K_longitudinal_ = 3 mM, K_longitudinal_+_lateral_ = 4 μM, weakening effect of GDP at the longitudinal interface = 50-fold, k_GTPase_ = 0.14 s^-1^

### Protein expression and purification

Plasmids to express wild type yeast αβ-tubulin have been described previously (Johnson *et al*., 2011; Ayaz *et al*., 2012) and were used without further modification. Plasmids encoding the β:C12A and β:C12S mutants were generated by site-directed mutagenesis. The integrity of constructs was confirmed by sequencing.

Wild type and mutant yeast αβ-tubulin were overexpressed in *S. cerevisiae* (Johnson *et al*., 2011). All proteins were purified using Ni-affinity and anion exchange chromatography as described previously (Johnson *et al*., 2011; Ayaz *et al*., 2012), dialyzed into storage buffer (10 mM PIPES pH 6.9, 1 mM MgSO4, 1 mM EGTA) containing 50 μM GTP (for polymerization measurements) or 50 μM GDP (for binding assays).

### Binding assays

Wild type or mutant yeast αβ-tubulin in 50 μM GDP was taken from −80 deg. C, rapidly thawed, and filtered (0.1 μm centrifugal filter, Millipore) at 4 deg. C to remove aggregates. To remove free GDP and GDP bound to αβ-tubulin, filtrate was exchanged into binding buffer (10 mM PIPES, 1 mM EGTA, and 1 mM MgSO_4_) using a Nick Sephadex G-50 gravity desalting column. A control experiment wherein ion exchange chromatography was used to quantify the nucleotide content of thusly treated tubulin showed that this procedure removes ~90% of the exchangeable nucleotide (data not shown). The concentration of αβ-tubulin in the filtrate was measured by UV absorbance using an extinction coefficient of 106835 M^-1^cm^-1^ (assuming all of the exchangeable GDP was removed).

To measure the affinity of S^6^-GTP for αβ-tubulin, we prepared 220 μl samples containing either 0.16 μM wild type or mutant αβ-tubulin in binding buffer with 0.5% Tween-20 and a variable concentration of S^6^-GTP. We used a competition assay to measure the affinity of GDP or dGDP for αβ-tubulin: samples were prepared as above but with 5 μM S^6^-GTP, and variable concentrations of competing nucleotide (GDP or dGDP) was added. All samples were kept in Eppendorf Protein LoBind tubes and allowed to equilibrate to RT before measurement.

Measurements of fluorescence quenching by S^6^-GTP and unquenching by GDP or dGDP were made in 96 well flat bottom black polystyrene plates using a Thermo Scientific Varioskan FLASH plate fluorimeter. 200 μl of each sample (including a blank) were loaded per well. Tryptophan fluorescence from αβ-tubulin was excited at 296 nm using a 12 nm slit width. Emission was monitored at 329 nm, with dynamic range set to medium high and a measurement time of 80 ms. Each sample was measured 100 times and mean fluorescence intensities were calculated. Replicate experiments were repeated multiple times on different days.

Because S^6^-GTP absorbs strongly at tryptophan’s emission peak, and both GDP and dGDP absorb at the excitation wavelength, it was necessary to correct our experiments for inner filter effects (Fishback and Yarbrough, 1984). To accomplish this we measured fluorescence from control samples consisting of ~50 μg/ml BSA in the assay buffer and a variable concentration of S^6^-GTP or GDP (GDP and dGDP have the same UV absorbance). The concentration of BSA was chosen to give about the same fluorescence intensity as 0.16 μM αβ-tubulin. Each sample was measured as described above, and mean fluorescence and standard error values were calculated. An exponential decay function fit to the mean fluorescence intensity values of each titration series (S^6^-GTP or GDP) and used to calculate correction factors that were later applied to the raw intensities recorded in quenching and unquenching experiments. For S^6^-GTP quenching experiments, we corrected fluorescence emission data using the expression F_corrected_ = F_observed_*EM_correction_([S6-GTP]). ‘EM_correction_’ denotes the inner filter correction for emission determined at different concentrations of S6-GTP. For GDP or dGDP competition experiments we corrected fluorescence emission data using the expression F_corrected_ = F_observed_*EM_correction_(5 μM S6-GTP)*EX_correction_([GDP]), where ‘EX_correction_’ is the inner filter correction for excitation determined at various concentrations of GDP.

Direct binding measurements were fit to a single-site saturation binding curve accounting for ligand depletion, and weighted by the standard errors (GraphPad Prism version 6). Competition binding measurements were fit to a single-site competition binding model, weighted by standard errors (GraphPad Prism version 6).

### GTPase activity assay

Wild type or mutant yeast αβ-tubulin in 50 μM GTP was taken from −80 °C, rapidly thawed, and filtered (0.1 μm centrifugal filter, Millipore) at 4 °C to remove aggregates. The concentration of αβ-tubulin in the filtrate was measured by UV absorbance using an extinction coefficient of 115000 M^-1^cm^-1^ to account for two bound guanine nucleotides. 1 μM αβ-tubulin of each species was polymerized in assembly buffer (100 μM K·PIPES, 2 mM MgSO_4_, 1 mM EGTA, pH 6.9) containing 1 mM GTP, 66 nM α-P32-GTP, 2 μM epothilone, and 10% glycerol at 30 °C. After 1 hr the reactions were centrifuged at 128,000 x g and 30 °C for 30 min. Pellets were washed 4X with assembly buffer to remove free nucleotide and unpolymerized αβ-tubulin. Pelleted MTs were denatured in 6 M Gua-HCl before being analyzed for nucleotide content by TLC. Nucleotide content was visualized by exposing X-ray film to the TLC plate.

### Time-lapse measurements of microtubule dynamics

Flow chambers were prepared as described previously (Gell *et al*., 2010). Sea urchin axonemes (Waterman-Storer, 2001) were adsorbed directly to treated coverglass before the blocking step to provide seeds for microtubule growth.

For dynamics assays, wild type or mutant yeast αβ-tubulin in 50 μM GTP was thawed, filtered and measured for concentration as described above. Protein was kept on wet ice for no more than 30 minutes before use in a MT dynamics assay. MT dynamics reactions were imaged by differential interference contrast microscopy (DIC) using an Olympus IX81microscope with a Plan Apo N 60x/1.42 NA objective lens and DIC prisms with illumination at 550 nm. Temperature was maintained at 30 deg. C by a WeatherStation temperature controller fit to the microscope’s body. The microscope was controlled by Micro-Manager 1.4.16 (Edelstein *et al*., 2010) and images recorded with a ORCA-Flash2.8 CMOS camera (Hamamatsu). Images were recorded every 500 ms for 1 to 2 hrs. At the end of each movie, a set of out-of-focus background images was taken for background subtraction (see below). To improve signal to noise, batches of 10 raw images were averaged using ImageJ (Schneider *et al*., 2012). The averaged images were opened as a stack and their intensities were normalized to the average image intensity before background subtraction. MT length was measured manually using the PointPicker plugin for ImageJ. Rates of MT elongation and catastrophe frequencies were determined as described previously (Walker *et al*., 1988).

### Dynamic Assays with Bim1-GFP

Samples containing either wild-type or β:C12A αβ-tubulin in the presence of 1 mM GTP or 2’dGTP, along with 50 nM Bim1-GFP in imaging buffer (BRB80 + 0.1 mg/mL BSA + 0.1% methylcellulose + antifade reagents (glucose, glucose oxidase, catalase), without the addition of β-mercaptoethanol (Gell *et al*., 2010)) were introduced into the flow chambers and imaged at 30 °C by total internal reflection fluorescence microscopy as described previously (Geyer *et al*., 2015). Images of MTs were taken every 5 seconds for 15-20 minutes. Bim1-GFP fluorescence intensity along microtubules and extending beyond their growing ends was obtained using the PlotProfile function in ImageJ (Schneider *et al*., 2012). The peak position of Bim1-GFP fluorescence was determined by fitting a gaussian to the intensity profile, and the line scans were shifted to align the peaks to the nearest pixel. The background intensity (as determined from the fluorescence in the line scan beyond the end of the MT) was subtracted before averaging the individual profiles.

**Supplemental Fig. 1.**
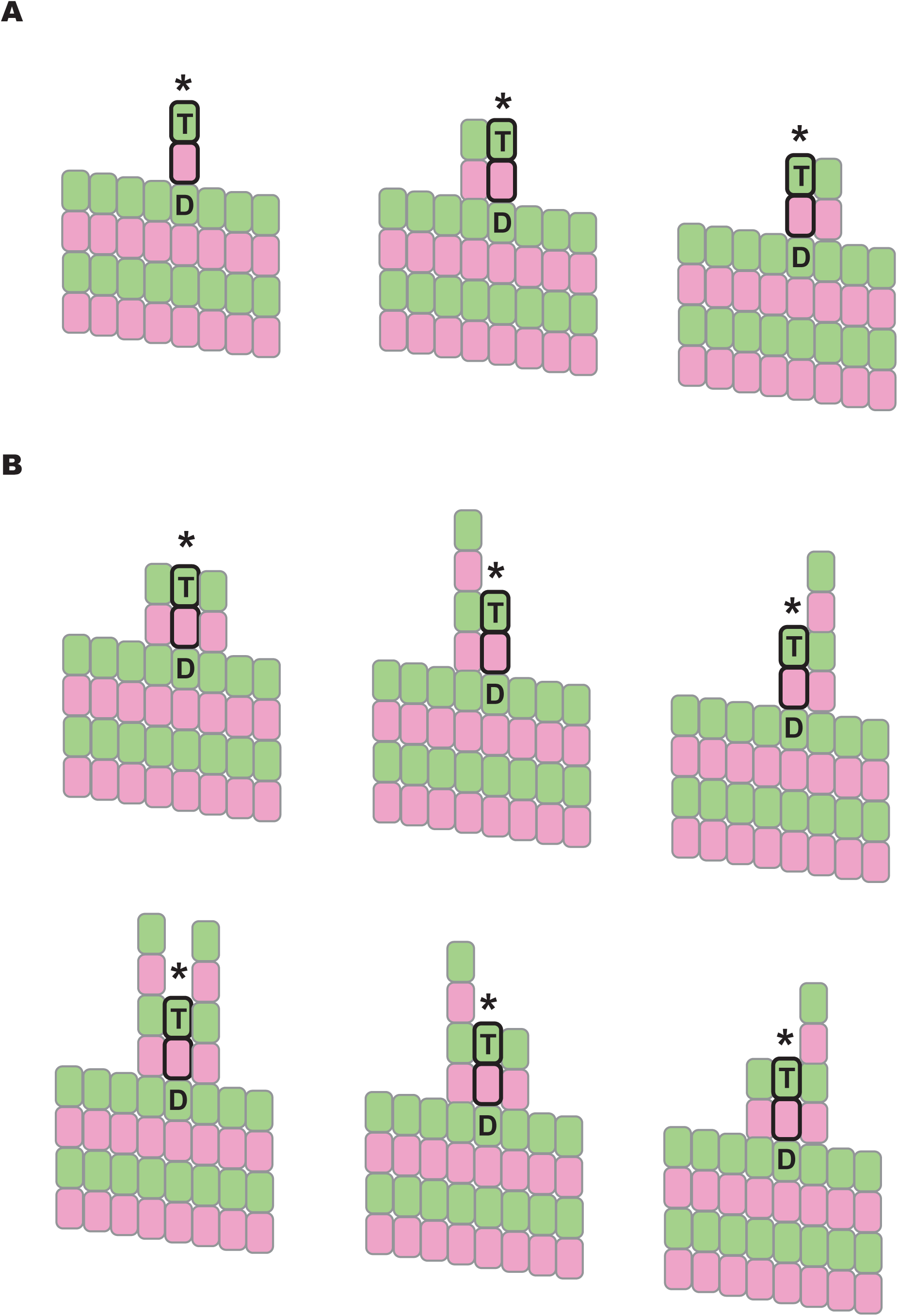
Elements of trans-action in an earlier model incorporating cis-acting GTP. (A) In the model from VanBuren et al., terminal GTP-bound subunits (bold and indicated by a *) that are above a GDP-bound subunit and that have zero or one neighbor as indicated were ‘tagged’ to dissociate according to the GDP rule even though that model assumed cis-acting GTP. This tagging was an attempt to implicitly capture mechanochemical features of the microtubule end (such as protofilament peeling) that were not modeled explicitly. (B) In that same model, terminal GTP-bound subunits that are above a GDP-bound subunit and that have two or more neighbors as indicated dissociated according to the GTP rule (cis action). We implemented these same assumptions in our tests of a cis model.

